# Young chicks rely on perceptual grouping to discriminate prime numbers

**DOI:** 10.1101/2021.03.04.433923

**Authors:** Maria Loconsole, Massimo De Agrò, Lucia Regolin

## Abstract

Grouping sets of elements into smaller, equal-sized, subsets constitutes a perceptual strategy employed by humans and other animals to enhance cognitive performance. We hypothesized that asymmetrical grouping, a characteristic of prime numbers, could provide visual cues enabling discrimination of prime from non-prime numerosities. Newborn chicks were habituated to even numerosities (as sets of elements presented on a screen), and then tested for their spontaneous choice among a prime and a non-prime odd numerosity. Chicks discriminated and preferred the prime over the composite set of elements. We discuss this result in terms of novelty preference. By employing different contrasts (i.e., 7vs.9, 11vs.9, and 13vs.15) we investigated the limits of such mechanism showing that induced grouping positively affects chicks’ performance.

## Main text

Adult humans can capitalize on non-symbolic perceptual mechanisms (e.g., grouping) to solve a symbolic task (e.g. enumeration)^1,2^. We are faster at enumerating a set of elements when these are divided into same-sized subsets (symmetrical grouping). When elements are not presented as grouped, we can actively implement the grouping strategy, although performance worsens with larger sets. Moreover, independently of the set numerosity, we are slower in enumerating prime numbers, an effect that has been ascribed to the impossibility to group a prime numerosity into same-sized subsets (asymmetrical grouping)^1^. Disassembling a numerosity into same-size subsets (symmetrical grouping) can constitute a non-mathematical strategy allowing to discern prime and non-prime numerosities^3^. Here we investigate whether day-old domestic chicks could capitalize on such symmetrical grouping strategy to discriminate between prime and non-prime sets of elements. Animals possess an intuitive sense of number from the very early stages of life. Such non-symbolic numerical abilities are widespread across species and are considered the evolutionary foundations of more complex numerical reasoning^4–7^. Birds display non-symbolic numerical abilities akin to humans, and can also exploit analogous cognitive and perceptual strategies, suggesting similarities in the underlying mechanisms^8–10^. In particular, young chicks allow early testing of visually guided behaviours, and soon after birth they were shown to process object symmetry^11,12^, behave according to Gestalt principles^13^, respond to spatial and numerical information^14^ and benefit from induced grouping strategies^15^.

To test our hypothesis, we investigated prime vs. non-prime discrimination in newly hatched chicks. In Exp.1 chicks were tested with either the 7vs.9 or the 11vs.9 discrimination, aiming at assessing a possible role of numerical magnitude (i.e., the prime numerosity could be either the larger or the smaller in the comparison). Additionally, by employing a more complex discrimination (i.e., 13vs.15) we explored processing limits in the capability to discriminate (Exp.2) and whether passively induced grouping (Exp.3, i.e., elements presented as already chunked by colour) could help chicks overcome such limits.

Each subject underwent a 1h habituation to sets (**Fig.1**) containing an even number of elements. Each set depicted a different colour-shape-number combination (i.e., yellow/blue/red/green x squares/rectangles/triangles x 6/8/10/12). Sets were presented one at a time on a screen according to a random sequence.

**Fig. 1.**
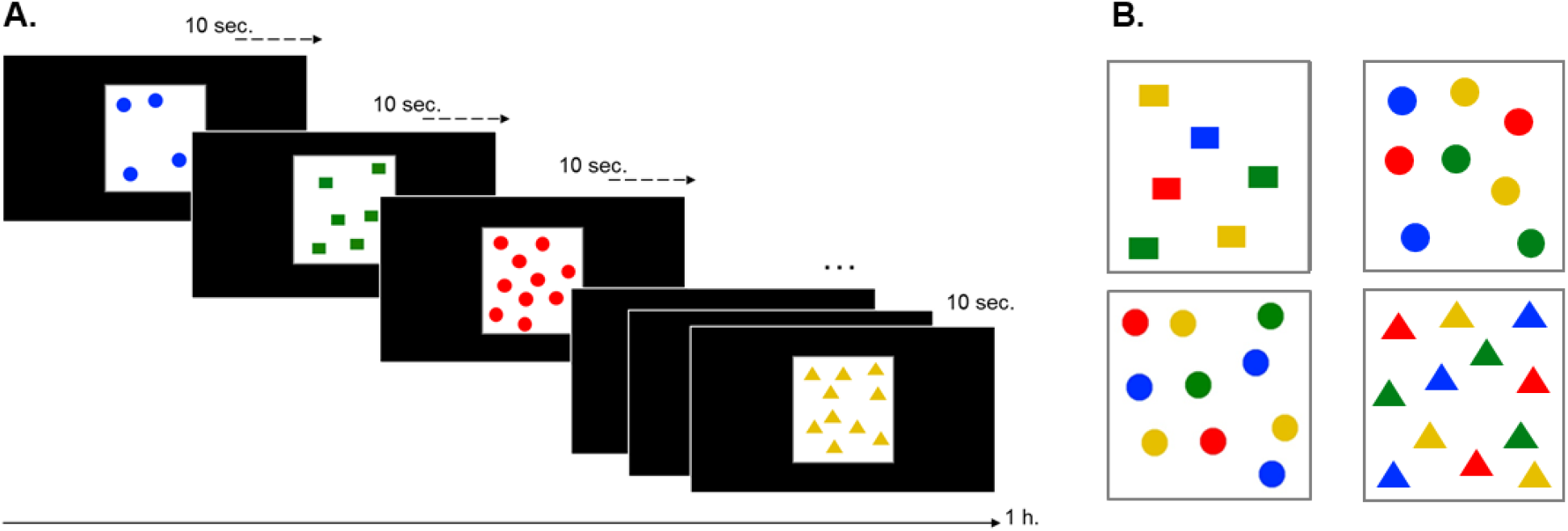
**A.** The habituation procedure (with examples of the stimuli used in Exp.1 and Exp.2). Each chick was individually exposed for 1h. to a sequence of stimuli depicting an even number of elements. Each stimulus appeared in the centre of the screen and after 10sec. it was replaced by a subsequent stimulus. In all experiments the sequence of the different stimuli was randomly determined. The position of the elements within the white area was pseudo-randomly determined so that two elements could never overlap. In Exps 1 and 2 the elements within each stimulus were of the same colour and shape. In Exp.3, elements within each stimulus were of four different colours (see B.) **B**. Examples of the multi-coloured stimuli used for habituation in Exp.3. Elements within the same stimulus had the same shape, whereas the colour of the elements was pseudo-randomly determined so that two elements located in close proximity from one another were never of the same colour.

At test (**Fig.2**), two sets appeared at the same time on two opposite sides of the arena (the same used for habituation). Sets simultaneously visible were always made of elements of the same colour and shape (i.e., taken from those used in the habituation) but depicted novel numerosities, one set being a prime and the other a non-prime odd number.

**Fig. 2.**
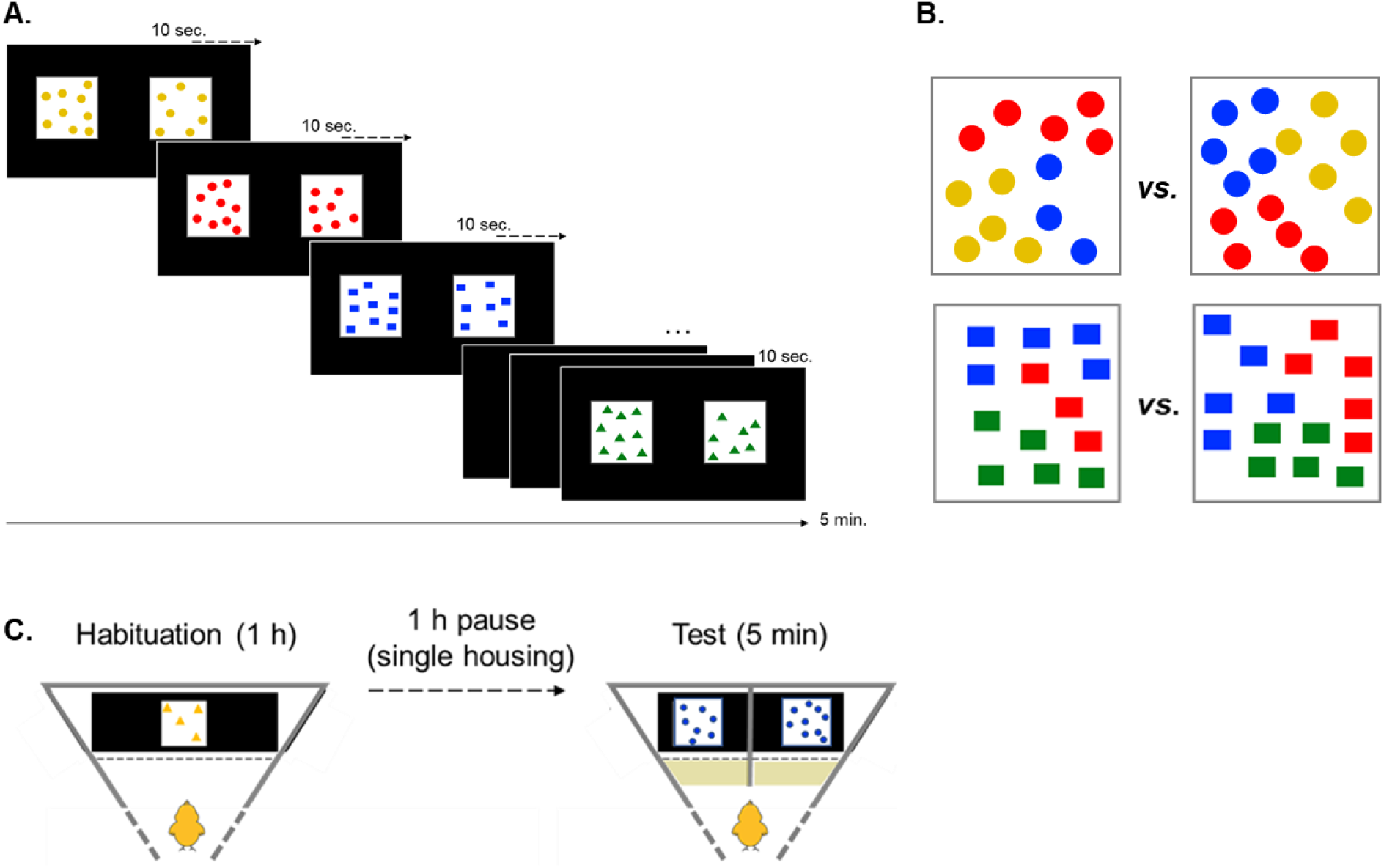
**A.** The test procedure (in the example, the 7vs.9 condition of Exp.1, with the prime numerosity presented on the right side). At test two stimuli were projected at once, one in the right and the other in the left half of the screen. Throughout the test the left-right position of the two numerosities remains identical for each subject (their position was randomized across subjects). During the 5 minutes of the test new pairs of stimuli were presented on the screen every 10sec. The two stimuli simultaneously presented were always of the same colour (except for Exp. 3, see B) and their elements were of the same shape. As for the habituation, the spatial arrangement of the elements in each stimulus was pseudo-randomly determined, and the presentation order of the stimuli followed a random sequence. **B**. An example of the testing stimuli used in Exp.3 (in this example the prime number is on the left side). Perceptual grouping was passively induced by presenting the elements as already chunked by colour into three subsets (same-coloured elements were always close to each other). In the stimulus depicting the prime numerosity (13) the three subgroups were made of 5, 5, and 3 elements; in the stimulus depicting the non-prime odd numerosity (15) each of the three subgroups comprised 5 elements. For the same chick the prime numerosity was presented in the same position (either left or right) throughout the test. The position of the prime numerosity was counterbalanced between subjects. **C**. Experimental procedure. During habituation (left) each chick saw for one hour a customized combination of multiple sets, each being individually presented on the screen for 10sec. At test (right) chicks were presented with pairs of stimuli comparing two odd numerosities (one prime and one composite). We scored the time each chick spent in either choice area (the shaded area in front of each stimulus) and considered it as a preference for the corresponding stimulus.

## Results

In Exp.1 (**Fig. 3A**) chicks were randomly assigned to the 7vs.9 (n=40) or to the 9vs.11 (n=39) comparison. We found an effect of number (GLMM analysis of deviance, *X*^2^=6.49; p=0.011): chicks spent longer by the prime (i.e., 7 or 11) rather than the composite (i.e., 9) numerosity in both comparisons (i.e., their preference was independent of numerical magnitude) (post-hoc analysis, estimate=-30.8; SE=11.8; t=-2.616; p=0.0098). The first approach was at chance (prob=0.557; SE=0.0561; z=1.004; p=0.351). In Exp.2 (**Fig. 3B**) we aimed to investigate whether chicks’ performance is affected by set size and tested them with 13vs.15 (n=39). In this case no significant difference emerged in the time spent near either stimulus (post-hoc analysis, estimate=31.6; SE=22.4; t=1.412; p=0.162) nor in the first set approached (post-hoc analysis, prob=0.475; SE=0.079; z=-0.316; p=0.752). In Exp.3 (**Fig. 3B**) grouping was passively induced as elements were chunked by colour into 3 smaller subgroups: 5+5+5 (symmetrical, for 15) and 5+5+3 (for 13) (Fig 1B). (n=40). In this case we found a preference for the prime numerosity both in the first approach (post-hoc analysis, prob=0.706; SE=0.078; z=2.326; p=0.02) and in the time spent by each stimulus, which was higher for 13 over 15 (post-hoc analysis, estimate=-83.5; SE=22.5; t=-3.707; p<0.001).

**Fig. 3.**
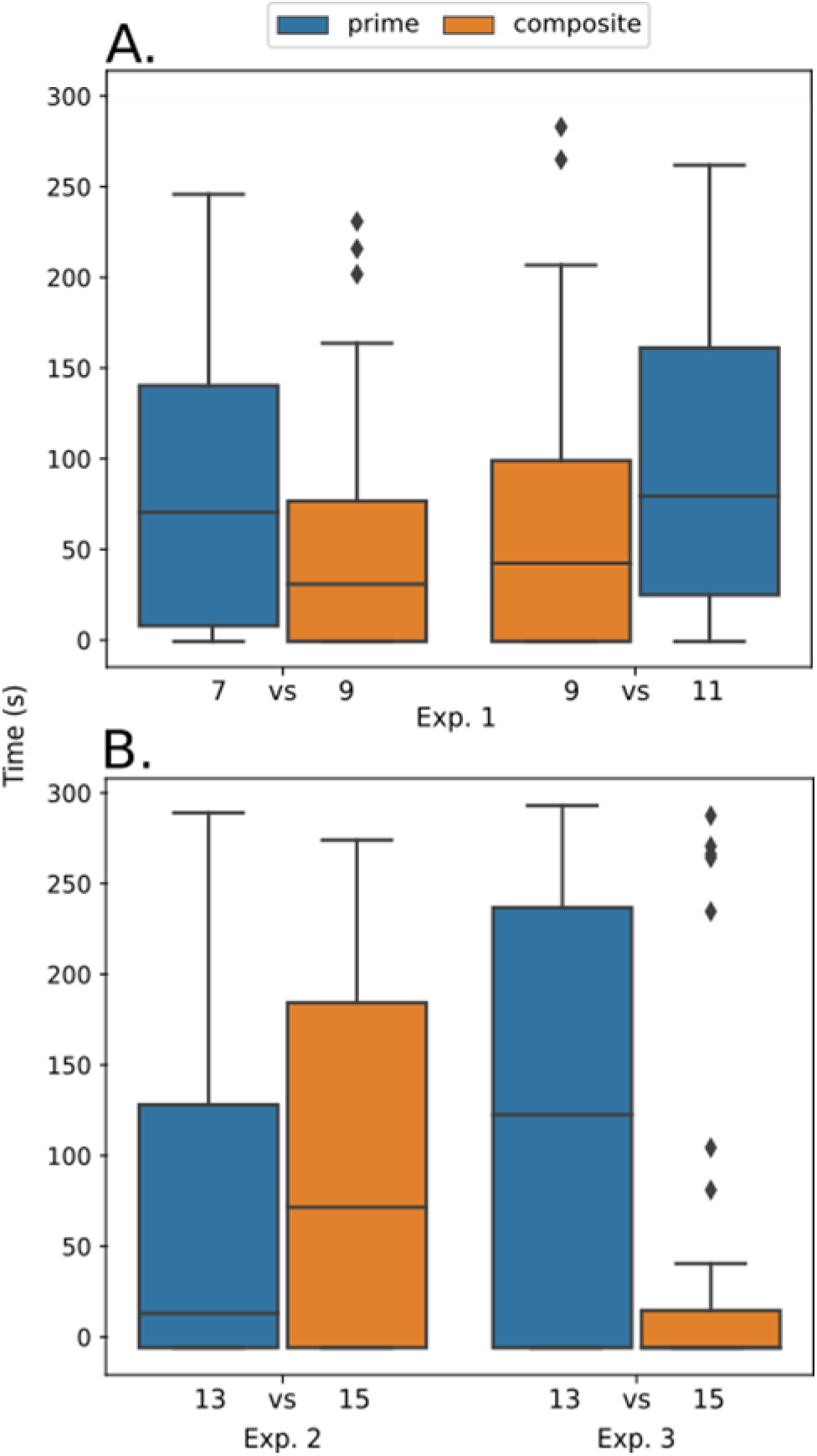
Time (sec) spent at test closer to the prime (blue) or to the composite (orange) numerosity. **A**. Exp.1: Chicks preferred the prime numerosity (7 or 11) irrespective of numerical magnitude. **B**. 13vs.15 comparison: in Exp.2 no preference for either stimulus emerged; in Exp. 3 (with induced grouping) chicks preferred the numerosity 13.

## Discussion

In the first experiment, chicks spent longer closer to the set of 7 or 11 elements rather than that of 9 elements. We interpreted these results as a preference for the prime numerosity. Avoidance of numerosity 9 has in fact never been reported, and in a previous study chicks did choose 9 in the 6vs.9 comparison^16^. Number-based strategies unlikely account for this result, nor can they explain direction of preference (choice of prime irrespective of it being larger or smaller in the comparison). We hypothesized discrimination relied on purely perceptual mechanisms such as disassembling the sets and comparing the resulting subgroups. Direction of the choice instead may have resulted from chicks’ preference for novelty and/or asymmetry, a well-known phenomenon in this species^11,17^. Prior to test chicks had been familiarized to even numerosities, which are more similar to non-prime odd numerosities. Both even and non-prime odd numerosities allow symmetrical grouping. E.g., 6 can be symmetrically grouped as 3+3, or 2+2+2; similarly, the odd non-prime number 9 can be disassembled in 3+3+3, also symmetrical. On the contrary, a prime number can only be grouped asymmetrically, i.e., at least one of the subsets comprises a different numerosity. The familiar test environment and familiar appearance of stimuli for colour and shape, likely prompted a preference for slight novelty (asymmetrical grouping)^17,18^. Early predispositions for visual asymmetry might have also played a role. These were reported for newly-hatched^11^ and week-old^12^ chicks and bear significant ecological value.

In the second experiment, we tested chicks in the 13vs.15 comparison. In human subjects, set numerosity affects performance: we are slower at enumerating a set as its numerosity increases^1,2^. We reported a similar effect in chicks, as they failed in discriminating between the two sets and chose randomly. We hypothesized a limit linked to working memory constraints in the number of subgroups (up to 3-4) and/or their size (up to 3-4 elements in each subgroup). Previous studies on numerical discrimination indicated a maximum limit of four “files” simultaneously represented in working memory^15^. Passively induced grouping (e.g. presenting the elements as already chunked) can help overcome such limit, enhancing performance in chicks^15^and infants^19^. If chicks relied on a symmetry-based perceptual mechanism, induced grouping should facilitate discrimination. Results from Exp.3 confirmed this hypothesis, showing that chicks could solve the discrimination, and that preference for the prime numerosity (i.e., 13) was restored. The emergence of a preference already in the first approach might constitute further evidence of induced grouping facilitating discrimination.

In conclusion, young chicks seem to represent and mentally manipulate sets of elements by disassembling them into smaller subgroups. As reported for adult humans^1,2^, this strategy implies symmetrical grouping and benefits from passively induced grouping^1,19^. These results provide the first experimental evidence of discrimination of prime numerosities in an animal model highlighting the presence of a spontaneous non-mathematical mechanism. This is in line with clinical evidence from individuals with a diagnosis of savant syndrome, who could recognize and/or generate prime numbers in the absence of mathematical skills^3,20^. Data from a non-mammalian species is particularly insightful as it implies an analogous and widespread mechanism in vertebrates, opening to the investigation of its neurobiological basis. Since our subjects were day-old, at least in this species mechanisms involved must be available very early during development and do not require formal training.

## Materials and Methods

### Subjects

We tested 158 domestic chickens (*Gallus gallus*). Fertilized eggs were provided by a local hatchery (Incubatoio La Pellegrina, San Pietro in Gu, PD, IT) and were incubated in the laboratory (Comparative Cognition Lab, Dept. of General Psychology, University of Padova) at controlled temperature (37.5°C) and humidity (55-66%) in a FIEM incubator MG 70/100 (cm 45 x 58 x 43). Soon after hatching, each chick underwent a 2h experimental procedure.

### Habituation

The apparatus consisted in two white plastic walls arranged to form a triangular arena (cm 93basis x 62length x 30height), with the longer side being a monitor (Samsung FHD, 24”, 60Hz) onto which the stimuli were projected. The vertex opposite to the monitor constituted the subject’s starting point (i.e., where the chick was gently placed at the beginning of the habituation). Habituation took place individually for each chick.

During habituation chicks were presented for 60 minutes with a random sequence of stimuli, each depicting an even number of elements (**Fig.1 A**). Every stimulus appeared in the center of the screen and remained visible for 10 seconds, then it was immediately replaced by the subsequent stimulus. Each stimulus was randomly selected to represent a combination of colour (i.e., red, blue, yellow, or green) and shape (i.e., triangles, rectangles, or circles) of its elements, as well as of the numerosity, always even, of the set (i.e., 4, 6, 10, or 12 elements). All elements comprised in a stimulus were positioned within a white squared area (336px) in the centre of the screen; the spatial location of each element was pseudo-randomly determined so that elements never overlapped with one another. Each element covered a total area of 36px.

In Exp. 1 and in Exp. 2, all the elements in a same stimulus were of the same colour and shape. In Exp. 3, elements within one stimulus had all the same shape, but they were of different colours so that all four colours available were presented within each stimulus. To avoid familiarizing the subjects with any sort of colour chunking during habituation, in Exp. 3 the elements located in close proximity were never of the same colour (**Fig.1 B**).

The habituation phase lasted 1 hour, during which the chick could freely move and approach the screen. The test took place 1 further hour after the end of habituation.

### Test

For Exp.1, day-old domestic chicks (N=79, 40♀) were asked to discriminate sets of either: 7vs.9 (N=40, 19♀) or 9vs.11 (N=39, 21♀) elements. In Exp.2 (N=39, 27♀) and Exp.3 (N=40, 19♀) chicks were tested in the discrimination of 13vs.15 elements.

For all experiments, the test arena was the same employed during habituation. But at test the screen was divided into two separate halves by a vertical plastic partition (cm 5basis x 30height). This way two test stimuli could be presented at once on the screen (i.e., either in the left and right half of it). The test consisted in a 5-minute presentation of pairs of novel stimuli (**Fig.2 A**). The two sets simultaneously displayed were identical for shape and colour of the elements but differed in their numerosity. In Exp.1 one stimulus was always a prime number, either 7 or 11, and was confronted with a stimulus depicting the composite numerosity 9 (symmetrical). In Exp.2 and Exp.3 a set of 13 elements (asymmetrical – prime) was confronted with a set made of 15 elements (symmetrical – composite). In all experiments the left-right position of the prime numerosity during the test was counterbalanced between subjects.

As for the habituation, random combinations of stimuli were presented throughout the test (a new pair of stimuli every 10 sec), the shape and colour of the elements randomly varied from pair to pair for the same chick.

In Exp.1 and Exp.2 all of the elements of the two stimuli simultaneously presented were of the same colour whereas in Exp.3 each stimulus comprised elements of three different colours. In this case same-colour elements were always adjacent to prompt grouping of the set into three subsets (i.e., 5+5+5 for the non-prime odd and 5+5+3 for the prime numerosity) (**Fig.2 B**).

Other than the shape-colour combination (the same 4 colours x 3 shapes used for the habituation), also the spatial disposition of the elements (within the white squared area) was randomly determined and differed for each individual stimulus and from trial to trial.

### Data analysis

The test was video recorded by a camera placed on top of the arena (Canon-Legria HF- R606). Video scoring was carried out offline and blind to the experimental hypothesis, and considered the time spent in each choice area, as well as the first stimulus approached by each subject. The data obtained were analyzed in R 3.6.3.

### Ethical Statement

The experiments complied with all applicable national and European laws concerning the use of animals in research and were approved by the Italian Ministry of Health (permit number: 196/2017-PR granted on 24/02/2017). All procedures employed in the experiments included in this study were examined and approved by the Animal Welfare Committee of the University of Padova (Organismo Preposto per il Benessere Animale, O.P.B.A.). At the end of the test all chicks were donated to local farmers.

## Supporting information

data analysis R code

raw data

## Acknowledgement

We wish to thank Ernesto Carafoli for discussing the original idea of the present study, Marco Bertamini, Jonathan N. Daisley and Peter Kramer for comments on earlier versions, Beatrice Guanciarossa and Lier Liu for chick care and data collection, Andreas Ioannis for data scoring. This work was carried out within the scope of the project “use-inspired basic research”, for which the Department of General Psychology of the University of Padova has been recognized “Department of Excellence 2018-2022” by the Italian Ministry of University and Research. M.L. is sponsored by a Ph.D. scholarship funded by the CARIPARO. Foundation.

## Author contributions

Maria Loconsole: Conceptualization, Methodology, Validation, Investigation, Writing - Original Draft. Massimo De Agrò: Methodology, Validation, Formal Analysis, Writing -Review &Editing. Lucia Regolin: Conceptualization, Methodology, Validation, Resources, Writing - Review &Editing, Supervision, Project Administration.

## Data and materials availability

Raw data and analysis code are available in the supplementary materials.

