## Supplementary material for "Young chicks rely on perceptual grouping to discriminate prime numbers": data analysis R code

**Full Analysis Description**

This supplement provides the entire R script and output of the statistical analysis we performed and the figures we produced, in their original form. It is presented in the spirit of open and transparent science but has not been edited in its style.
We scored the time (seconds) spent by each stimulus, within a minimum distance of 5cm from the screen.

Data preparation

```
load packages
library(emmeans) #for post-hoc
library(readxl) #to import data from excel
library (DHARMa) #to check goodness of fit

## This is DHARMa 0.3.2.0. For overview type '?DHARMa'. For recent changes, type
news(package = 'DHARMa') Note: Syntax of plotResiduals has changed in 0.3.0, see
?plotResiduals for details

library (car) #to do Anova on glm

## Loading required package: carData
## Registered S3 methods overwritten by 'car':
##   method                          from
##   influence.merMod                lme4
##   cooks.distance.influence.merMod lme4
##   dfbeta.influence.merMod         lme4
##   dfbetas.influence.merMod        lme4

library (glmmTMB) #to do mixed models
library(ggplot2) #to plot
```

Experiment 1

There are two conditions, one in which the prime number is the smaller of the two presented (7vs9) and one in which it is the larger (9vs11).

Exp.1 - Preliminary analysis

Preference for the biggest numerosity

First, we create the model.

```
pms <- glmmTMB(time~size+(1|subj),data = exp1, family = gaussian)
```

Testing goodness of fit

```
simres<-simulateResiduals(pms) #standard seed for random values is 123
plot(simres, asFactor=T)
```

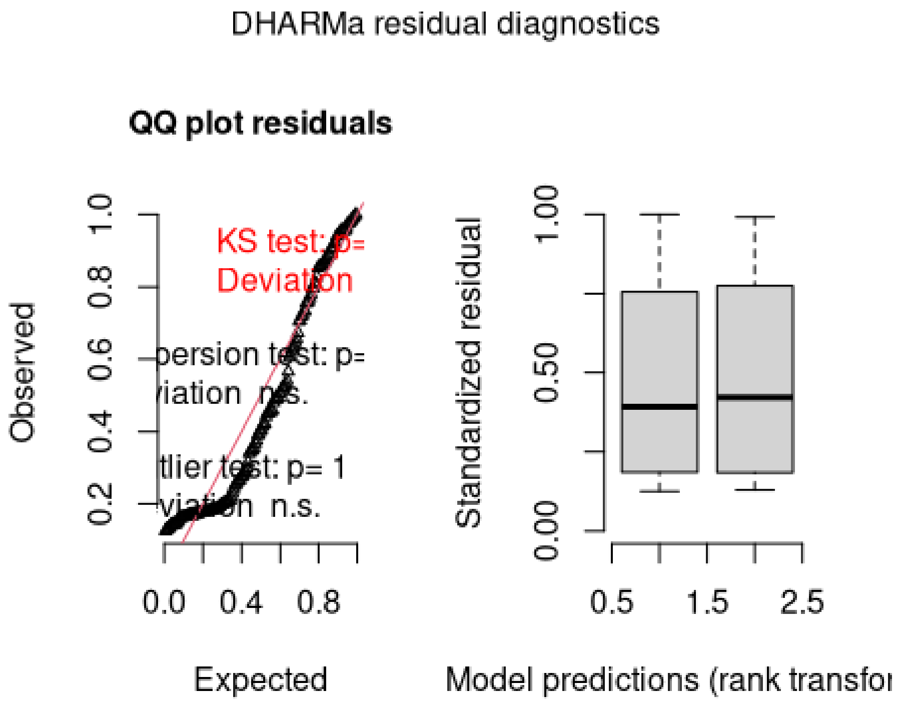


Deviation is significant. It looks like there is a lot of deviation in the low part, while the distribution is good for higher numbers. maybe lots of zeros?

```
hist(exp1$time,100)
```

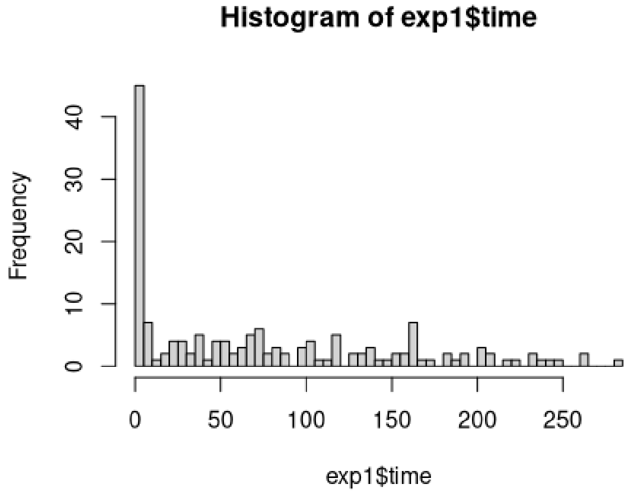


Yes, indeed. However, a correction for zero inflation may not be proper. Are these zero representing a total inactivity, or a very clear choice?

```
tapply(exp1$time,exp1$subj,sum)
```

```
## S001 S002 S003 S004 S005 S006 S007 S008 S009 S010 S011 S012 S013 S014
S015 S016
##  119   73  148  202   75  165  100   39  237   96  203  143   51  162  121   33
## S017 S018 S019 S020 S021 S022 S023 S024 S025 S026 S027 S028 S029 S030
S031 S032
##   85  246  222  191  161  255   49  205  185  101  186   74  216   95   25   66
## S033 S034 S035 S036 S037 S038 S039 S040 S041 S042 S043 S044 S045 S046
S047 S048
##  151  157  174   40  160  231  160  212  193  187  127  199  125  102   68  287
## S049 S050 S051 S052 S053 S054 S055 S056 S057 S058 S059 S060 S061 S062
S063 S064
##  205  156   80   81  156  243  165  118  135  161  133   71  283   95   74  171
## S065 S066 S067 S068 S069 S070 S071 S072 S073 S074 S075 S076 S077 S078
S079
##  111  137  193  227  205  161  207  164  124  262  184  265  117  161  179
```

Here printed the total number of second for each subject near either stimulus. No subject scored a total of zero seconds near either stimulus, meaning that any subject that spent 0 second near one of the two stimuli actually stayed near the other, meaning a clear and robust choice. Data is not zero inflated, zeros are actually an important value.
We will use the previously created model.

```
glmmTMB:::Anova.glmmTMB(pms)

## Analysis of Deviance Table (Type II Wald chisquare tests)
##
## Response: time
##       Chisq Df Pr(>Chisq)
## size 0.1514  1     0.6972
```

No effect of stimulus size, so there is no preference for either small or big stimuli in the experiment.

<u>Exp.1 - First choice</u>
We want to see if there is a difference in the first approached stimulus. We coded as 1 (correct) a first approach towards the prime, 0 (wrong) the composite. Model will not have random effect, as to each subject correspond a single row of the table. Also, we will not include the "stimulus type" as a factor, as the difference is already coded in the 1, 0 nature of the dependent variable

```
pmf <- glm(first~cond,data = exp1, family = binomial)
Anova(pmf)
## Analysis of Deviance Table (Type II tests)
##
## Response: first
##      LR Chisq Df Pr(>Chisq)
## cond  0.60894  1     0.4352
```

no effect of condition, just need to see if the choice is different than 50%

```
e<-emmeans(pmf,~1)
test(e, type="response")
##  1       prob     SE  df z.ratio p.value
##  overall 0.557 0.0561 Inf 1.004   0.3152
##
## Results are averaged over the levels of: cond
## Tests are performed on the logit scale
```

No preference

<u>Exp.1 - Main Analysis</u>

In the main analysis, we want to see if the animals have a preference for the prime numerosity. We will include in the model, other than condition and stimulus; the subjects' sex, as according to the literature it may have some effect

```
mm <- glmmTMB(time~stim*cond*sex+(1|subj),data = exp1, family = gaussian)
## Warning in fitTMB(TMBStruc): Model convergence problem; non-positive-definite
## Hessian matrix. See vignette('troubleshooting')
we are getting a convergence problem. to get more info
summary(mm)
##  Family: gaussian  ( identity )
## Formula:          time ~ stim * cond * sex + (1 | subj)
## Data: exp1
##
##      AIC      BIC   logLik deviance df.resid
##       NA       NA       NA       NA      148
##
## Random effects:
##
## Conditional model:
##  Groups   Name        Variance  Std.Dev.
##  subj     (Intercept) 1.115e-16 1.056e-08
##  Residual             5.440e+03 7.376e+01
## Number of obs: 158, groups:  subj, 79
##
## Dispersion estimate for gaussian family (sigma^2): 5.44e+03
##
## Conditional model:
##                      Estimate Std. Error z value Pr(>|z|)
## (Intercept)            65.646     16.921   3.879 0.000105 ***
## stimprime              25.767     23.931   1.077 0.281591
## cond9vs11               3.231     23.354   0.138 0.889962
## sexM                  -15.089     23.354  -0.646 0.518220
## stimprime:cond9vs11    -9.182     33.027  -0.278 0.781007
```

```
## stimprime:sexM            -1.669     33.027  -0.051 0.959700
## cond9vs11:sexM             3.221     33.267   0.097 0.922871
## stimprime:cond9vs11:sexM   41.693     47.047   0.886 0.375510
## ---
## Signif. codes:  0 '***' 0.001 '**' 0.01 '*' 0.05 '.' 0.1 ' ' 1
```

The random effect is 0, meaning there is no variance between subjects. in this case is normal to get this error. I could remove the random effect, but The results will be the same as this model.

```
glmmTMB:::Anova.glmmTMB(mm)
## Analysis of Deviance Table (Type II Wald chisquare tests)
##
## Response: time
##               Chisq Df Pr(>Chisq)
## stim          6.4954  1    0.01082 *
## cond          0.7973  1    0.37191
## sex           0.1193  1    0.72977
## stim:cond     0.2335  1    0.62896
## stim:sex      0.6442  1    0.42221
## cond:sex      1.0468  1    0.30625
## stim:cond:sex 0.7854  1    0.37551
## ---
## Signif. codes:  0 '***' 0.001 '**' 0.01 '*' 0.05 '.' 0.1 ' ' 1
```

We found only the effect of stim variable. This means:

- No difference between sexes, nor in general activity (sex variable) nor in condition, nor in choices (interactions)
- No difference between conditions. This means that any preference would not change between conditions (no interaction effect) and that conditions are not different in difficulty for subject (the total choice time is the same)
- An effect of the stimulus, meaning that chicks indeed prefer one between prime and composite numerosity. I will do a post-hoc to determine the direction of their preference.

```
e<-emmeans(mm,~stim) #leaving only stim as is the only effect found
## NOTE: Results may be misleading due to involvement in interactions
pairs(e)
##  contrast          estimate   SE  df t.ratio p.value
##  composite - prime    -30.8 11.8 148 -2.616  0.0098
##
## Results are averaged over the levels of: cond, sex
```

Chicks prefer the prime over the composite numerosity.

<u>Exp.1 – Plotting results</u>

```
ggplot(exp1,aes(x=cond,y=time,fill=stim))+ #nice!
```

geom_boxplot()

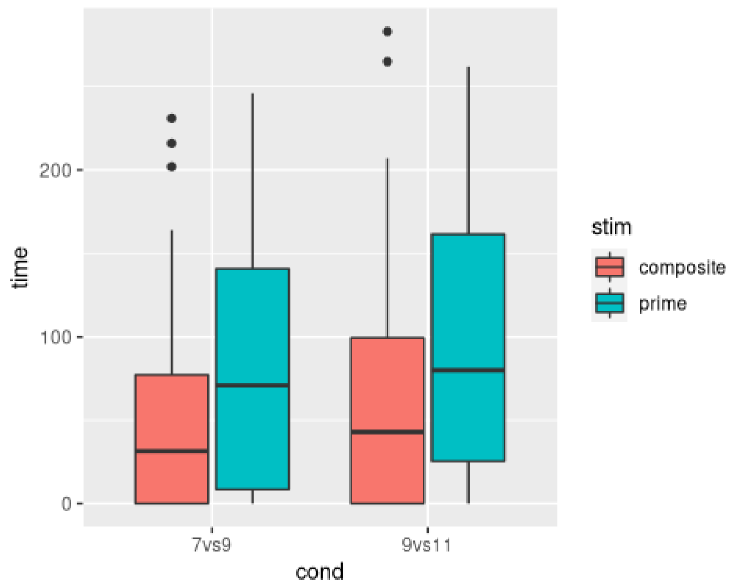


Experiment 2

In the first experiment, condition 9vs11 is surprising, as generally chicks have a hard time in distinguishing these two numerosities. We wanted to push this even forward, with the 13vs15 comparison.

Preliminary analysis

No preference for the larger numerosity to check here, as the prime is always the smallest. Directly to first choice

First choice

```
pmf2 <- glm(first~1,data = exp2, family = binomial)
```

No factors, so not doing anova. directly post-hoc

```
e<-emmeans(pmf2,~1)
test(e, type="response")
##  1       prob    SE  df z.ratio p.value
##  overall 0.475 0.079 Inf -0.316  0.7519
##
## Tests are performed on the logit scale
```

No preference

Main Analysis

We follow the same procedure as before

```
mm2 <- glmmTMB(time~stim*sex+(1|subj),data = exp2, family = gaussian)
```

```
simres<-simulateResiduals(mm2) #standard seed for random values is 123
plot(simres, asFactor=T)
```

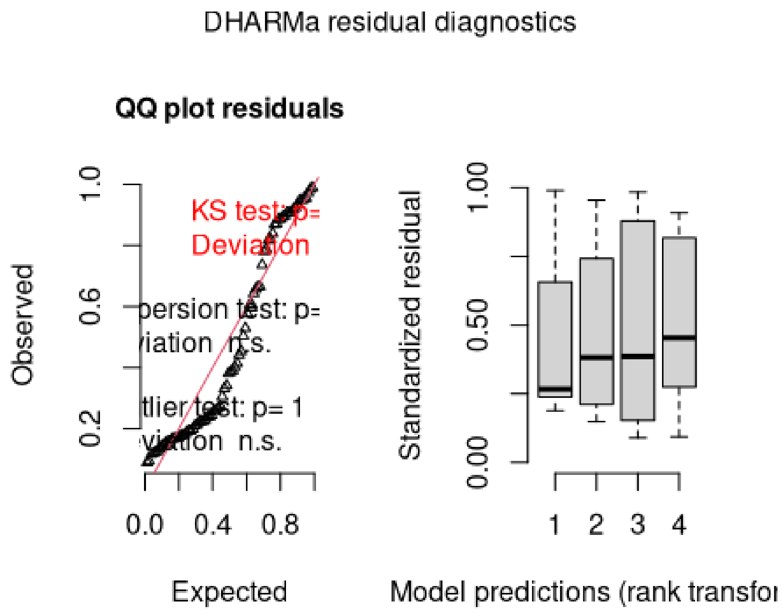


Still deviating, but seems different from before. check for zero inflation nontheless

```
hist(exp2$time,100)
```

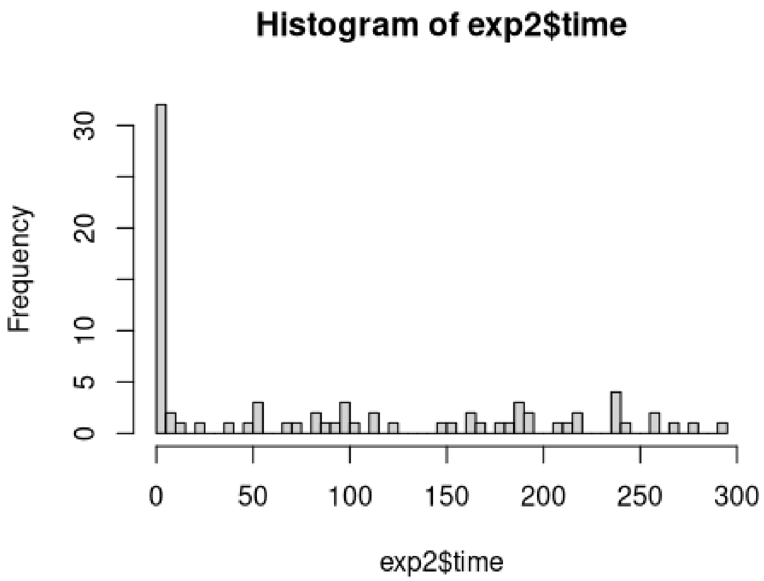


```
tapply(exp2$time,exp2$subj,sum)
```

```
## S080 S081 S082 S083 S084 S085 S086 S087 S088 S089 S090 S091 S092 S093
S094 S095
```

```
## 236 194 267 178 142  24 240 124 210 241 111 189 211 178  97 192
## S096 S097 S098 S099 S100 S101 S102 S103 S104 S105 S106 S107 S108 S109
S110 S111
## 187 295 216  93   8 149   7  83 185 238 218 223 164 189 256 256
## S112 S113 S114 S115 S116 S117 S118 S119
## 280 124 236 212 151 165  40 240
```

We are in the same situation as before, just with more distance between high and low performing chicks. As before we will keep the analysis on the same model.

```
glmmTMB:::Anova.glmmTMB(mm2)
## Analysis of Deviance Table (Type II Wald chisquare tests)
##
## Response: time
##           Chisq Df Pr(>Chisq)
## stim     1.7332  1     0.1880
## sex      0.0224  1     0.8809
## stim:sex 0.2620  1     0.6087
```

No effect emerges. We will do a post-hoc for completeness, to look at the raw data, but we have to accept the null hypotesis for both conditions in exp2

```
e<-emmeans(mm2, ~stim)
## NOTE: Results may be misleading due to involvement in interactions
pairs(e)
##  contrast          estimate   SE df t.ratio p.value
##  composite - prime     31.6 22.4 74 1.412   0.1620
##
## Results are averaged over the levels of: sex
e
##  stim      emmean   SE df lower.CL upper.CL
##  composite  104.5 15.8 74     73.0      136
##  prime       72.9 15.8 74     41.4      104
##
## Results are averaged over the levels of: sex
## Confidence level used: 0.95
```

Raw data may suggest a reversing effect (preference for larger numerosity), but it is not significant. Also, the SE is overall much higher.

<u>Exp.2 - Plotting</u>

```
ggplot(exp2,aes(x=stim,y=time))+
  geom_boxplot()
```

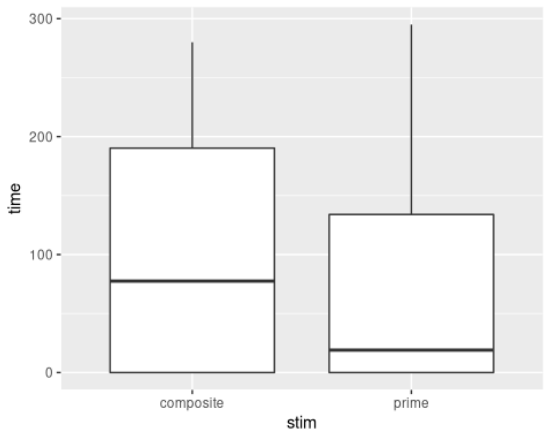


Note how both boxplot have the 0 in Q1. This is why there is no significant effect even if the average of composite number is higher: unlike experiment 1, different subjects have completely different preference.

Experiment 3 – Induced colour grouping

In Exp3, we wanted to test if we can support discrimination by passively inducing the gruping strategy (i.e., present the elements grouped by colour).

Experiment 3 – Preliminary analysis
No preference for biggest numerosity to check here, as the prime is always the smallest. Directly to first choice

Experiment 3 – First choice

```
pmf3 <- glm(first~1,data = exp3, family = binomial)
```

No factors, so not doing anova. directly post-hoc

```
e<-emmeans(pmf3,~1)
test(e, type="response")
##  1        prob     SE  df z.ratio p.value
##  overall 0.706 0.0781 Inf 2.326   0.0200
##
## Tests are performed on the logit scale
```

we already find a preference for the prime number at first choice (as 1 is for prime, 0 is for composite)

Experiment 3 – Main Analysis
We follow the same procedure as before

```
mm3 <- glmmTMB(time~stim*sex+(1|subj),data = exp3, family = gaussian)
```

```
simres<-simulateResiduals(mm3) #standard seed for random values is 123
plot(simres, asFactor=T)
```

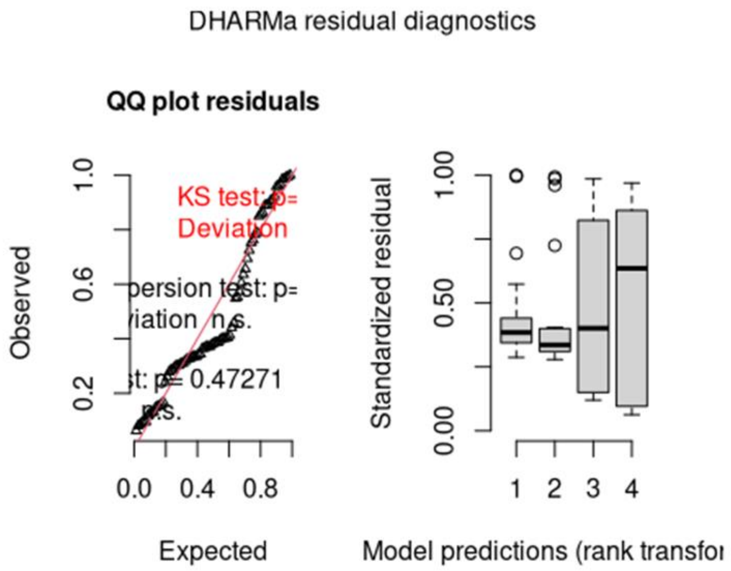


Still deviating.

```
hist(exp3$time,100)
```

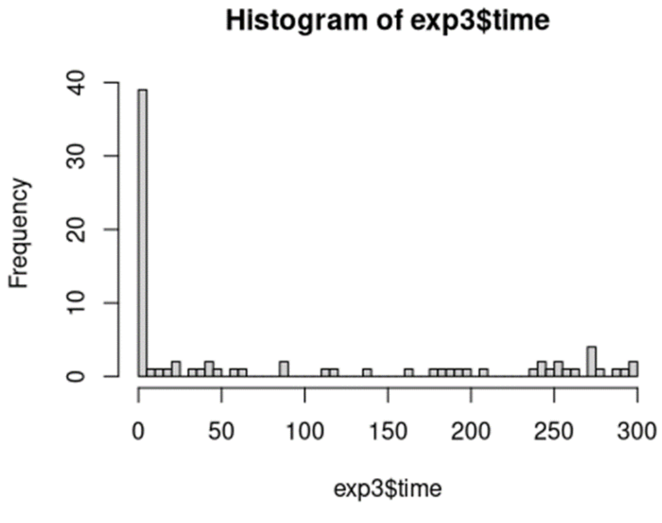


```
tapply(exp3$time,exp3$subj,sum)
##    S120    S121    S122    S123    S124    S125    S126    S127    S128    S129
## 263.718 274.850 276.545 178.025 255.204 235.465 105.118   0.000 200.018
63.737
##    S130    S131    S132    S133    S134    S135    S136    S137    S138    S139
## 285.260 212.730 206.706 118.724   0.000  12.996  17.402 270.157 240.575 272.137
```

```
##    S140    S141    S142    S143    S144    S145    S146    S147    S148    S149
## 296.765 240.625 252.302 123.291 241.842 273.687   0.000 253.339   0.000 293.409
##    S150    S151    S152    S153    S154    S155    S156    S157    S158    S159
## 197.646 249.206  51.473 163.968   0.000 299.013 138.279 180.220  44.990   0.000
```

Again, a lot of zeros. In this situation there are some chicks that shown no preference at all, but only 4 out of 40. We will still proceed with the model, following the same considerations as before.

```
glmmTMB:::Anova.glmmTMB(mm3)
## Analysis of Deviance Table (Type II Wald chisquare tests)
##
## Response: time
##            Chisq Df Pr(>Chisq)
## stim     14.1177  1  0.0001717 ***
## sex       0.1455  1  0.7028817
## stim:sex  0.8225  1  0.3644498
## ---
## Signif. codes:  0 '***' 0.001 '**' 0.01 '*' 0.05 '.' 0.1 ' ' 1
```

No effect of sex, but there is an effect of the stimulus (prime or composite). Not surprising given the binomial result.

```
e<-emmeans(mm3, ~stim)
## NOTE: Results may be misleading due to involvement in interactions
pairs(e)
##  contrast          estimate   SE df t.ratio p.value
##  composite - prime    -83.5 22.5 74 -3.707  0.0004
##
## Results are averaged over the levels of: sex
e
##  stim      emmean   SE df lower.CL upper.CL
##  composite   42.9 15.9 74     11.2     74.6
##  prime      126.4 15.9 74     94.7    158.1
##
## Results are averaged over the levels of: sex
## Confidence level used: 0.95
```

We see a strong preference for the prime number

<u>Exp.3 - Plotting</u>

```
ggplot(exp3,aes(x=stim,y=time))+
  geom_boxplot()
```

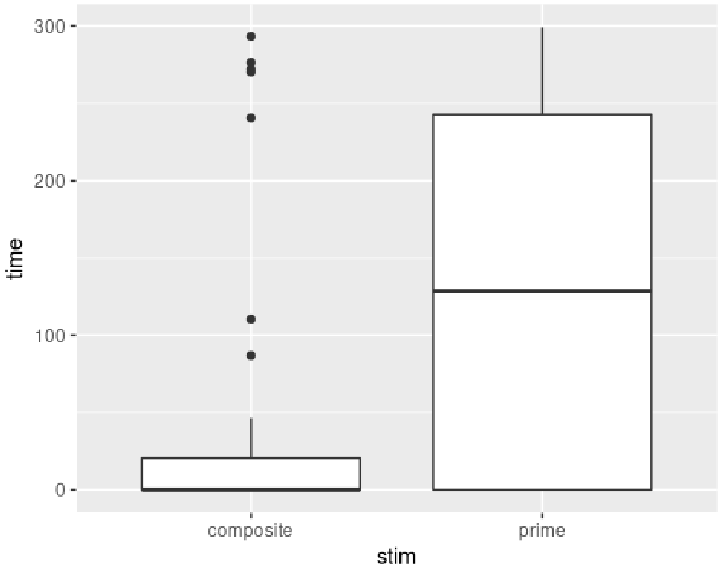
300
200
100
0
time
composite
prime
stim
